# Sensory plasticity in a socially plastic bee

**DOI:** 10.1101/2022.01.29.478030

**Authors:** Rebecca A Boulton, Jeremy Field

## Abstract

The social Hymenoptera have contributed much to our understanding of the evolution of sensory systems. Attention has focussed chiefly on how sociality and sensory systems have evolved together. In the Hymenoptera, the antennal sensilla are important for optimising the perception of olfactory social information. Social species have denser antennal sensilla than solitary species, which is thought to enhance social cohesion through nest-mate recognition. In the current study, we test whether sensilla numbers vary between populations of the socially plastic sweat bee *Halictus rubicundus* from regions that vary in climate and the degree to which sociality is expressed. We found region level differences in both olfactory and hygro/thermoreceptive sensilla numbers. We also found evidence that olfactory sensilla density is developmentally plastic: when we transplanted bees from Scotland to the south-east of England, their offspring (which developed in the south) had more olfactory hairs than the transplanted individuals themselves (which developed in Scotland). The transplanted bees displayed a mix of social (a queen plus workers) and solitary nesting, but neither individual nor nest phenotype was related to sensilla density. We suggest that this general, rather than caste-specific sensory plasticity provides a flexible means to optimise sensory perception according to the most pressing demands of the environment. Sensory plasticity may support social plasticity in *H. rubicundus* but does not appear to be causally related to it.

## Introduction

Natural selection on sensory abilities has resulted in enormous diversity in the mechanisms that organisms use to perceive their environment. For example, species of Lake Tanganyika cichlids that have colonised more complex, rocky habitats have greater visual acuity than sand-dwelling species (Dobberfuhl et al. 2005). These mechanisms can also feedback into selective regimes, facilitating micro- and macroevolutionary change through sensory drive. Sensory drive occurs when local adaptation results in populations with diverging modes of communication and perception. This has been proposed to lead to assortative mating between partners with complementary signalling and perception systems, reducing geneflow and promoting speciation (Boughman 2002).

The Hymenoptera have emerged as a model system to understand both the proximate and ultimate factors that underlie sensory evolution. These insects have antennal receptors with diverse functions and distinct forms that can easily be characterised within and between species. This work has revealed that both the density and form of antennal sense organs (sensilla) correlate with diet (Polidori et al. 2012; 2020), sex (Babu 2019; Do Carmo Queiroz Fialho et al. 2014) and a cleptoparasitic lifestyle (Wcislo 1995; Galvani et al. 2017). Attention has focussed particularly on how eusociality has imposed selection to optimise antennal perception. Eusociality involves reproductive division of labour with a queen/s producing eggs and workers provisioning the brood. However, sociality takes diverse forms across the bees and wasps and group sizes vary enormously, providing rich opportunities for comparative study. Many obligately social species show worker polyphenism, where some workers specialise on brood care and others on foraging or guarding the nest (Andersson 1984). Compared with solitary species, social species require more finely tuned communication systems to distinguish intruders from kin and gain information from nestmates (Wcislo 1997; Renner and Nieh 2008; Freeberg et al., 2012; Peckre et al., 2019). Across the Hymenoptera, social species tend to have greater densities of antennal sensilla that are involved in the perception of olfactory cues than solitary species (Elgar et al. 2018). This is thought to enhance social cohesion and performance by facilitating nest-mate recognition (Ozaki et al. 2005; Gill et al. 2013; Couto et al. 2017) and supporting caste polyphenisms (Spaethe et al. 2007; Riveros and Gronenberg 2010; Gill et al. 2013; Grüter et al. 2017; Elgar et al. 2018) without requiring the production of costly and complex chemical signals (Kather & Martin 2015).

The olfactory antennal sensilla have been proposed as a pre-adaptation that acts alongside haplodiploidy to predispose the Hymenoptera to the evolution of sociality (Couto et al. 2017). A conserved, ancestral olfactory subsystem, involving basiconic antennal sensilla that detect chemical information from nestmates and neural circuits that process it, has been suggested to facilitate the evolution of sociality in the Hymenoptera. This works by allowing nestmate recognition from cuticular hydrocarbon (CHCs) profiles (Ozaki et al. 2005; McKenzie et al. 2016; Couto et al. 2017; Pask et al. 2017 see also Kather et al., 2016). While empirical evidence for this subsystem exists for only two species (an ant and a hornet), the halictid bees have provided support for the possibility that sensory pre-adaptations might contribute to the evolution of sociality. The Halictidae or sweat bees exhibit a range of social structures, from solitary nesting through to obligate sociality, including within-species social polymorphism and social plasticity (Gibbs et al. 2012). A comparative study by Wittwer et al (2017) has shown that ancestrally solitary species and their social sister species have equivalent densities of hair-like sensilla (basiconic and trichoid sensilla) that detect olfactory cues. Species that have reverted to a solitary existence from sociality, on the other hand, have reduced densities of these sensilla. This suggests that in the Halictidae, sensitivity to olfactory cues precedes the evolution of sociality rather than evolving as a consequence of it.

In addition to caste polyphenisms, where individuals always form social groups but vary in the tasks that they perform, a number of species are facultatively social. In these species, individuals vary in their propensity to form social groups or nest as solitary individuals (Gibbs et al. 2012). This social plasticity is thought to be key to the evolutionary diversification of the Hymenoptera as it provides a genetic base from which obligate eusociality and caste polyphenism can evolve repeatedly (West-Eberhard 2003; Rehan & Toth, 2015; Jones et al. 2017; Shell & Rehan, 2018). It also provides an opportunity to test for correlates of social evolution without the complication of interspecific differences (see also Schradin 2013; Schradin et al., 2018). In this study we characterise the antennal sensilla of a facultatively social species, *Halictus rubicundus*, for the first time. This species shows regional differences in social behaviour which are not fixed and depend on the climate (see methods below; Field et al., 2010; 2012). Phylogenetic work suggests that social plasticity in *H. rubicundus* has evolved from an ancestral state of obligate sociality in the genus *Halictus* (Danforth 2002).

We measure the density of different antennal sensilla types between populations which vary in their expression of sociality due to the climate and test whether sensory plasticity mirrors social plasticity in a transplant experiment with *H. rubicundus*. Our results represent an important step in disentangling the sequence of events that leads to the evolution of eusociality and reproductive division of labour, as well as the evolutionary loss of these traits in the Hymenoptera.

## Methods

### Study system

*Halictus rubicundus* is a ground-nesting bee that can be found throughout Europe and North America in areas with sandy-loamy soil that receive sufficient sunlight. In Great Britain and Ireland, *H. rubicundus* demonstrates social variation according to climate (latitude and altitude) but retains social plasticity throughout its range (Field et al. 2010; 2012). In the north and at high altitudes bees are typically solitary as the short growing season and cooler climate precludes the bivoltinism that is required for social living. In a typical year in colder climates, foundresses produce a single brood of offspring per year, the females of which enter hibernation and emerge to become foundresses the following spring (figure 1a). In areas further south and at low altitudes growing seasons tend to be longer and the weather is warmer, so bivoltinism and sociality can occur more frequently. After overwintering underground, females produce a first brood (the B1) which become workers and provision a second brood (B2) which are offspring of either the original foundress or a B1 replacement queen. B1 females are also observed to provision their nests alone and lay eggs (without workers, i.e. become solitary foundresses) in the same year that they emerge. The B2 hibernate and emerge the following year to found their own nests (figure 1b; Field et al. 2010; 2012).

**Figure 1 :**
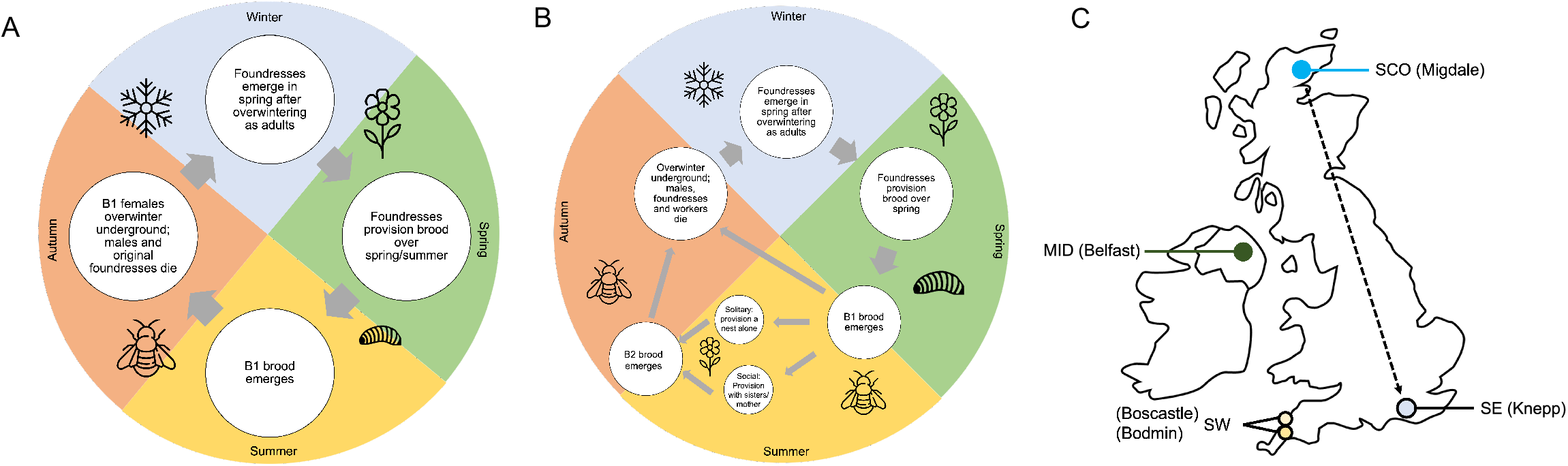
(A) univoltine lifecycle of *Halictus rubicundus* which prohibits social nesting (B) bivoltine lifecycle of *Halictus rubicundus* where sociality is possible as the 1st brood (B1) can help to provision a second brood (B2). Note that in (B) the B1 females can either (i) overwinter and provision the following year as a solitary foundress (ii) provision a nest alone as a solitary foundress in the year that they emerge or (iii) provision the nest socially with their B1 sisters with a queen (their mother or a sister) in the year that they emerge. (C) Map depicting the sites where *H. rubicundus* females were studied. Dashed arrow represents the direction of the transplant experiment (SCO to SE).

Sociality is extremely plastic in *H. rubicundus* in Great Britain, and when solitary northern bees are transplanted to the south, the B1 brood will often provision as workers or solitary foundresses in the same year (Field et al. 2010; 2012; this study). Here we test for fixed and plastic differences in antennal sensilla density in *H. rubicundus* across and within populations. We first test whether olfactory sensilla densities differ between bees from southern populations, where the climate allows some individuals to form social nests most years, and northern populations with limited opportunity to form social nests. Second, we test whether antennal sensilla numbers are developmentally plastic by transplanting bees from the north to the south. Do the offspring of northern bees that develop in the warmer south have different numbers of antennal sensilla than their parental generation which developed in the north? We chose to conduct a north-to-south transplant because we wanted to test for differences in sensilla number between individual and nest-level phenotypes, to determine whether social individuals/nests exhibit greater densities of olfactory sensilla to aid in nestmate recognition. This comparison is possible only when bees were able to express their full range of social behaviour (i.e. in the warmer south).

### Collection

*Halictus rubicundus* females were collected from 4 sites in the UK in 2018, 2019 and 2020 (see fig 1c). Two sites in the south-west (SW): Boscastle (Cornwall: N 50° 41’ 24” W 4° 41’ 24”; N = 12; 2018) and Bodmin (Cornwall, N 50° 30’ 36” W 4° 33’ 36”; N = 4; 2019); a site at a mid-latitude (MID): Belfast (Northern Ireland, N 54° 32’ 24” W 5° 58’ 48”; N = 19; 2020); and a site in the far north of Scotland (SCO): Migdale (Scotland, N 57° 53’ 24” W 4° 15’ 0”; N = 15; 2018 and 2020). Previous studies and our own observations indicate that while a proportion of nests are social in SW (this study), nests are always non-social in Belfast (MID; Field et al. 2010) and Scotland (SCO; this study). However, bees from Belfast (MID; Field et al. 2010) and Scotland (SCO; this study) are socially plastic and may have social nests when moved to more southerly sites. Bees sampled were expected to be foundresses based on collection date. We collected females by hand-netting in late June in SCO, and in May and early June in the SW, when all bees had worn wings indicative of several weeks of provisioning. In Belfast (MID) we excavated overwintering foundresses in February.

### Transplant

In addition to sampling from native populations, we collected bees that had been transplanted from Scotland (SCO) to a site in the south-east (SE), the Knepp estate in West Sussex (N 50° 53’ 60” W 0° 21’ 36”; N = 47; see fig 1c). We refer to these transplanted bees as ‘T’ bees hereafter. Buckets of soil were embedded within a nest aggregation at the Scottish site during winter 2018-19, and native foundresses subsequently nested in them during 2019. Their offspring (T) then emerged in late summer 2019 and hibernated in the buckets. The buckets containing hibernating T bees were taken from Scotland to the south-east in the spring of 2020, where they were re-embedded in the ground. The T foundresses were marked with a dot of enamel paint when they provisioned a B1 brood in the south-east. When the B1 emerged, they were marked with a different colour of enamel paint. Each nest was marked with a numbered nail at the entrance. This allowed us to distinguish fresh B1 females from the original foundresses and to determine whether nests were solitary (with a single B1 female or original foundress provisioning alone) or social (with multiple B1 females provisioning the nest). At the end of the B1 provisioning phase we collected T foundresses (which had emerged in Scotland and been transplanted to the SE) and their B1 offspring (which had developed and emerged in the SE; we refer to these as ‘B1’ individuals).

### Specimen storage and preparation

All specimens were stored in 95% Ethanol until they were prepared for imaging. One antenna was removed from each bee (whether it was the left or right was recorded) and mounted on a JEOL aluminium stub (10mm dia × 10mm high) using a PELCO carbon conductive sticky tab (10mm dia). Specimens were mounted dorsal side up. Specimens were carbon coated using spectrograpically pure carbon.

### Imaging

Mounted and carbon coated specimens were imaged using a TESCAN Vega SEM in High Vacuum mode at 20.0kV. The scan mode was set to resolution and the magnification was 485x. Only the two distal segments (11 and 12), which have the highest density of sensilla, were imaged. The length and area of each antennal segment was measured during imaging and images of each antennal segment from the SEM were saved in TIF format.

### Image scoring

Eight distinct types of sensilla with overlapping functions have been characterised in the Hymenoptera (Do Carmo Queiroz Fialho et al. 2014). Placoid sensilla are plate-like structures that are involved in olfaction and chemoreception. Tricoid and basiconic sensilla are hair-like projections with multiple sub-types that are used in olfaction involving contact. Coeloconic, campaniform and ampulliform sensilla are pore-like and are involved in sensing temperature, humidity and CO2 concentration (Do Carmo Queiroz Fialho et al. 2014). For this study we grouped all sensilla types into three structural/functional groups that allowed for repeatability in scoring: (i) olfactory plate sensilla (sensilla placodea, fig 2), (ii) olfactory hair-type sensilla (sensilla trichodea and basoniconica; fig 2) and (iii) temperature/humidity/CO2 pore sensilla (coeloconic/campaniform/ampulliform sensilla; fig 2).

**Figure 2 :**
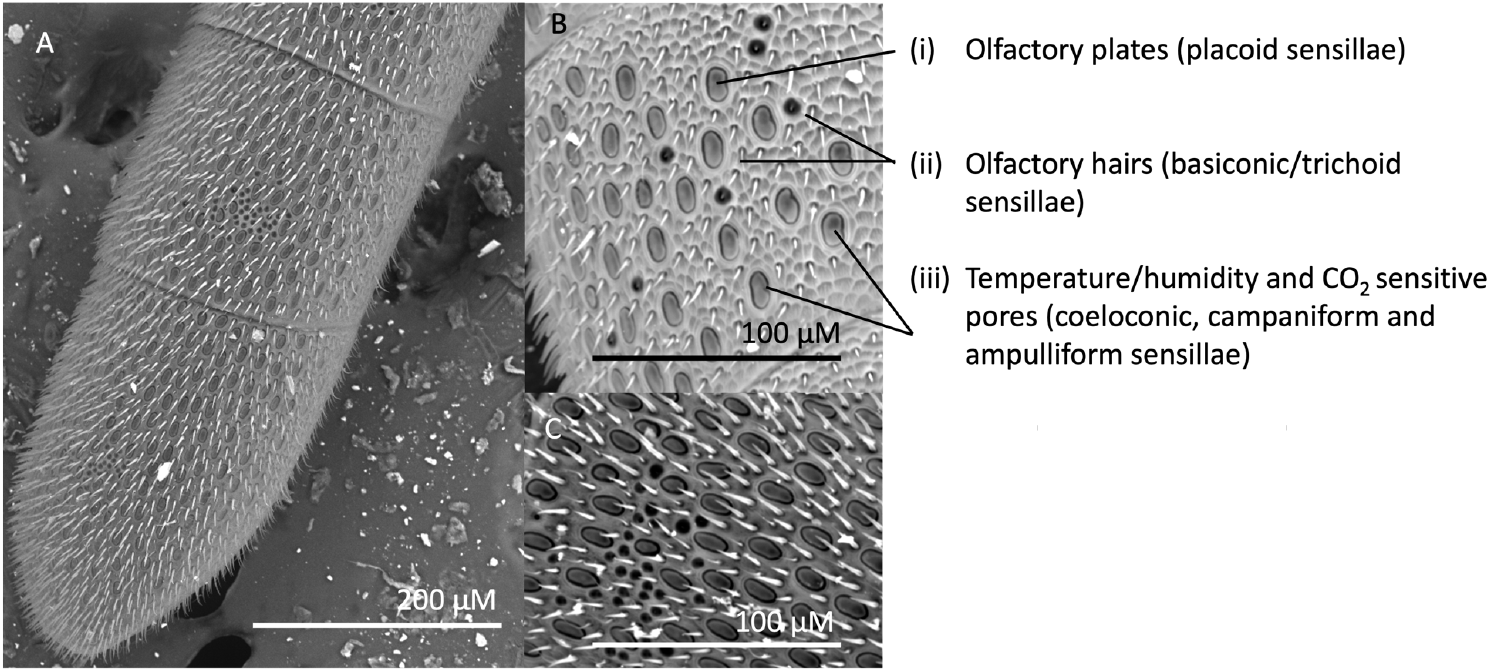
SEM image showing a *Halictus rubicundus* antenna tip. (A) shows the distal 2 antennal segments (11 and 12) that were imaged, (B) and (C) show the types (i, ii, and iii) of antennal sensilla that were counted. Scale is given in µM.

Images of antennal segments 11 and 12 were scored using ImageJ software. For each segment, three 50µM x 50µM ‘quadrats’ were selected, the X and Y coordinates of which were derived by generating a random number using the excel function RANDBETWEEN. Each quadrat was saved as a TIF image and the number of sensilla types i and ii were counted. We counted sensilla type iii across the entire segment as they are distributed unevenly on the surface of the antenna. In some cases, bare patches with no sensilla were present on the surface of the antenna. Any quadrats which fell on the bare areas were discarded and another area was selected. We note that the scorer was not blind to the population of origin as the same person was required to mount, image and score the antennae.

### Analysis

#### Repeatability

Thirty-three of the 130 sampled antennae were scored a second time (by the same person) for all sensilla types. The repeatability of image scoring was assessed using the intra-class correlation coefficient, function ICC in the R package irr (Gamer et al, 2019).

#### Sensilla numbers

We used linear mixed models in the R package lme4 (Bates et al. 2019) to test for differences in the numbers of sensilla types (i-iii). For all models outlined below, the response variable was total sensilla counted, either i) olfactory plates, total across 3 quadrats for each segment per individual; ii) olfactory hairs, total across 3 quadrats for each segment per individual; or iii) temperature/humidity pores, total across each segment per individual. We consider the unit of replication (N) to be a single bee. We did not measure more than one antenna from each bee, but we did measure 2 segments per antenna. As such, Individual was fitted as a random effect in all models. The package DHARMa was used to determine the most appropriate error structure for all models (Gaussian) and to interpret residual plots for lack of fit (Hartig, 2020).

#### Region, development site and social phenotype

To test whether bees from different regions have different numbers of sensilla we included region as a fixed factor with 3 levels in LMMs: SW (Boscastle and Bodmin combined; N = 16 individuals), MID (Belfast, Northern Ireland; N = 19 individuals) and SCO (Migdale, Scotland, far north; N = 15 individuals).

We ran additional models using only data from bees in the transplant experiment to test whether sensilla density (type i, ii or iii) varies depending on where bees completed their development (which may suggest developmental plasticity). To do this we compared bees that developed in Scotland (‘T’ foundresses) with their offspring that developed in the south-east (B1 females) after transplantation. LMMs included a fixed effect of development site, with 2 levels: SCO (bees that emerged in Scotland and were either collected there (N = 15 individuals, same individuals that were used in the region models; or ‘T’ females that were moved to the south-east as adults, N = 4) and SE (bees that emerged as adults after immature development in the south-east, the B1 offspring of the bees that were transplanted from Scotland to the south-east; N = 27). Note that *H. rubicundus* females mate before overwintering and so both the mother and father of the B1 bees collected in the south-east originated in Scotland.

Additional LMMs were used to test whether B1 offspring collected in the SE (using the same individuals used in the development site model for which nest phenotypes were known, N = 14) had higher counts of sensilla if they were workers (hereafter B1w) from social nests of transplanted females compared to those which left and started their own solitary nests (hereafter B1sol). Social nests had a queen and one or more workers (B1w), while solitary nests had a single B1sol foundress that produced offspring in the same year. For these models we included nest phenotype as a fixed effect with 2 levels (social, N = 6 or solitary, N = 8). We did not sample multiple individuals from the same nest, so did not include nest as a random effect. We did not recover any queens from social nests and so could not test for an effect of bee phenotype within social nests. We also ran a model to test for effects of individual, rather than nest-level, phenotype across social and solitary nests, using the same bees. For this analysis bees were characterised as (1) a future reproductive, B1hib (a B1 individual that had emerged in 2020 and did not provision that year, will hibernate and provision in 2021; N = 13); (2) a worker, B1w (a B1 individual that provisioned a nest containing a queen; N = 7); (3) a solitary foundress from 2019, T (a foundress that emerged in SCO in 2019, was transplanted to the south-east and laid eggs/provisioned a nest alone without workers); (4) a solitary foundress from 2020, B1sol (a foundress that emerged in the B1 generation in 2020 in the SE and provisioned a nest alone; N = 4).

Finally, we ran a model using the bees collected in the SE to test whether age-related wear and tear reduces density of all 3 sensilla types. For these analyses age was binary: bees were scored as fresh (newly emerged and had not provisioned a nest; N = 12) or old (had provisioned a nest for several weeks N = 16).

All LMMs included antenna segment number (11 or 12) as a fixed factor. The interaction effect between region/development type/social phenotype and segment number was also fitted. We included a random effect of individual as two segments were imaged for each bee. Models were run using the function lmer in the R package lme4 (Bates et al, 2015).

## Results

### Repeatability, differences between antennal segments and effects of age

Counts of all sensilla types were highly repeatable (Koo and Li 2016; ICC greater than 0.85 for all; for a detailed summary see supplementary material table S1). Olfactory plates (type i) and olfactory hairs (type ii) were found in greater numbers on the most distal antennal segment, segment 12, while hygro/thermoreceptors (type iii) were found in higher numbers on segment 11.

We also found no evidence that age-related wear and tear influences the variation in sensilla density; there was no effect of age on density of any of the three types of sensilla (type i: X2 = 0.25, df = 1, p = 0.62; type ii: X2 = 0.01, df = 1, p = 0.92; type iii: X2 = 0.09, df = 1, p = 0.76).

## Region

There was a significant effect of region on the counts of type ii (olfactory hairs) and iii (thermo/hygroreceptors) sensilla, but not type i (olfactory plate receptors; see table 1, figure 3a-c). Pairwise tests show that bees from mid-latitude (MID; Belfast; N = 19) had significantly more olfactory hairs (type ii) than bees collected from the far north (Scotland; N=30; p = 0.03; figure 3b), whereas bees from the south-west (N = 16) did not differ from MID (N = 32; p = 0.06) or Scottish (N = 15; p = 0.93) bees. Bees from SCO and MID had more thermo/hygro (type iii) receptors than SW bees (p < 0.05; figure 3c). Bees from the south-west had the most olfactory plate sensilla (type i) but this was not statistically significant (figure 3a). There were no interaction effects between segment number and geographic region (table 1).

**Table 1 :** LMM results showing effect of region on antennal sensilla counts across segments using type II sums of squares (p-values for main effects are calculated independent from interaction effects which in all cases are non-significant).

| Sensilla type | Effect | $X^2$ | df | P |
| --- | --- | --- | --- | --- |
| Type i) olfactory plates | Region | 2.31 | 2 | 0.31 |
| | Segment | 19.48 | 1 | $1.01 \times 10^{-5}$ |
|  | Region*Segment | 0.45 | 2 | 0.80 |
| Type ii) olfactory hairs | Region | 7.81 | 2 | 0.02 |
| | Segment | 41.74 | 1 | $1.04 \times 10^{-10}$ |
|  | Region*Segment | 2.70 | 2 | 0.26 |
| Type iii) thermo/hygro-receptive | Region | 9.14 | 2 | 0.01 |
| | Segment | 19.17 | 1 | $1.20 \times 10^{-5}$ |
|  | Region*Segment | 0.35 | 2 | 0.84 |

**Figure 3 :**
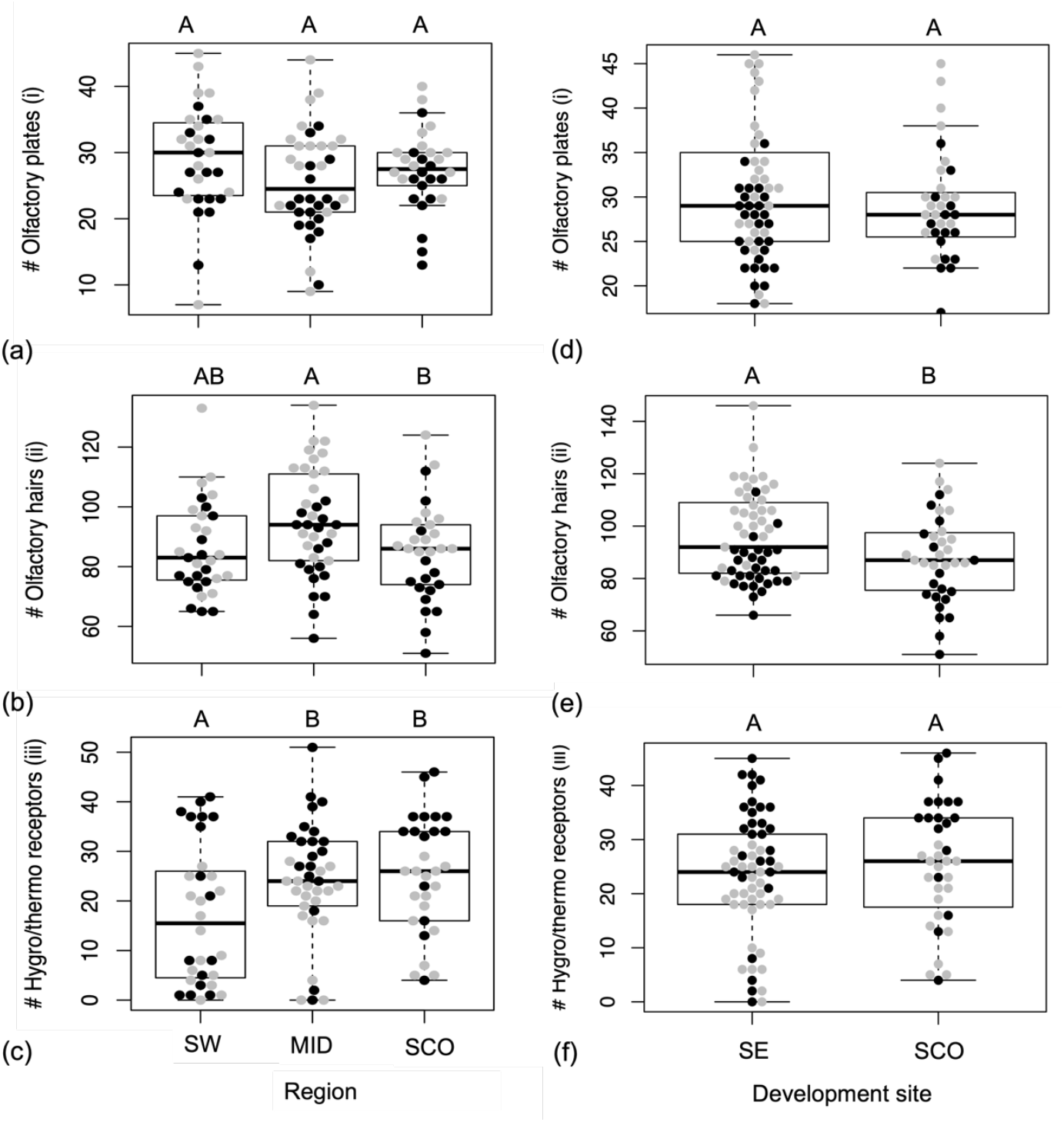
Numbers of (a) and (d) olfactory plates, i (top row); (b) and (e) olfactory hairs, ii (middle row); and (c) and (f) hygro/thermoreceptive pores, iii (bottom row), for Halictus rubicundus across: a-c three sampled regions: SW (South-west; mostly social), MID (Belfast; solitary) and SCO (Scotland; solitary); d-e for Halictus rubicundus originating from Scotland and collected after either developing as larvae in Scotland (SCO) or after developing as larvae at the transplanted site in the south-east (SE). Different uppercase letters (A or B) over bars indicate significantly different counts of sensilla. Black points represent sensilla counts on segment 11 and grey points on segment 12. Box plots show median and quartiles across segments 11 and 12.

### Development site

The density of olfactory hairs (type ii) varied according to where bees originating from Scotland spent their immature development (table 2; figure 3d-e). Offspring (B1) of transplanted bees (T) which developed in the south-east (N=27) had more olfactory hairs (type ii, p = 0.01) than conspecifics that developed in their natal Scottish site (N=19; fig 3e). A similar trend was seen in olfactory plates (I; fig 3d) but this was not statistically significant (p = 0.06). There was no effect of developmental site on thermo/hygroreceptor numbers (iii; fig 3f).

**Table 2 :** LMM results showing effect of developmental site on antennal sensilla counts across segments (p-values for main effects are calculated independent from interaction effects which in all cases are non-significant).

| Sensilla type | Effect | X<sup>2</sup> | df | P |
| --- | --- | --- | --- | --- |
| Type i) olfactory plates | Development site | 3.44 | 1 | 0.06 |
|  | Segment | 34.82 | 1 | 3.6x10<sup>-9</sup> |
|  | Development site *Segment | 0.61 | 1 | 0.43 |
| Type ii) olfactory hairs | Development site | 6.41 | 1 | 0.01 |
|  | Segment | 61.35 | 1 | 4.79x10<sup>-15</sup> |
|  | Development site *Segment | 1.57 | 1 | 0.21 |
| Type iii) thermo/hygroreceptive | Development site | 0.74 | 1 | 0.39 |
|  | Segment | 33.25 | 1 | 8.12x10<sup>-9</sup> |
|  | Development site *Segment | 0.15 | 1 | 0.69 |

### Sociality

B1 workers (B1w) from social nests (N= 6) and solitary B1 (B1sol) females that founded new nests alone (N= 8) had equivalent numbers of all 3 sensilla types (i-iii, see table 3; fig 4a-c) and there were no interactions between nest phenotype and antenna segment number. There was some suggestion that solitary bees had higher numbers of olfactory plates (type i), but this was not statistically significant (p = 0.07). Similarly there were no differences in numbers of any sensilla types across different individual phenotypes (B1hib future reproductive emerged in 2020 in SE and did not provision that year, will hibernate and provision in 2021, N = 13; B1w worker emerged in 2020 in the SE, N = 7; T solitary foundress that emerged 2019 in SCO and provisioned in the SE in 2020, N = 4; and B1sol solitary foundress emerged and provisioned in 2020 in SE, N = 4; see table 4; figure 4d-e). These results were the same if B1 solitary foundresses from 2019 (that developed and emerged in SCO) were excluded (see archived code and data; https://github.com/DrBecky-B/Bee.Antennae).

**Table 3 :**
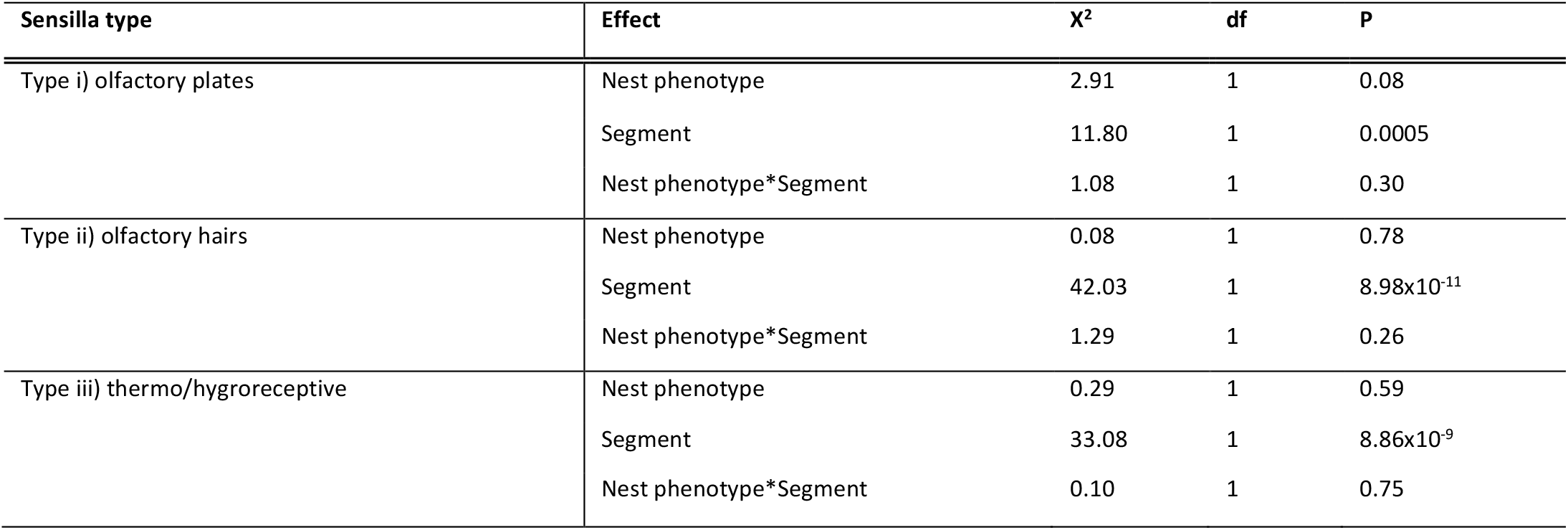
LMM results showing effect of nest phenotype (social/solitary) on antennal sensilla counts across segments (p-values for main effects are calculated independent from interaction effects which in all cases are non-significant).

**Figure 4 :**
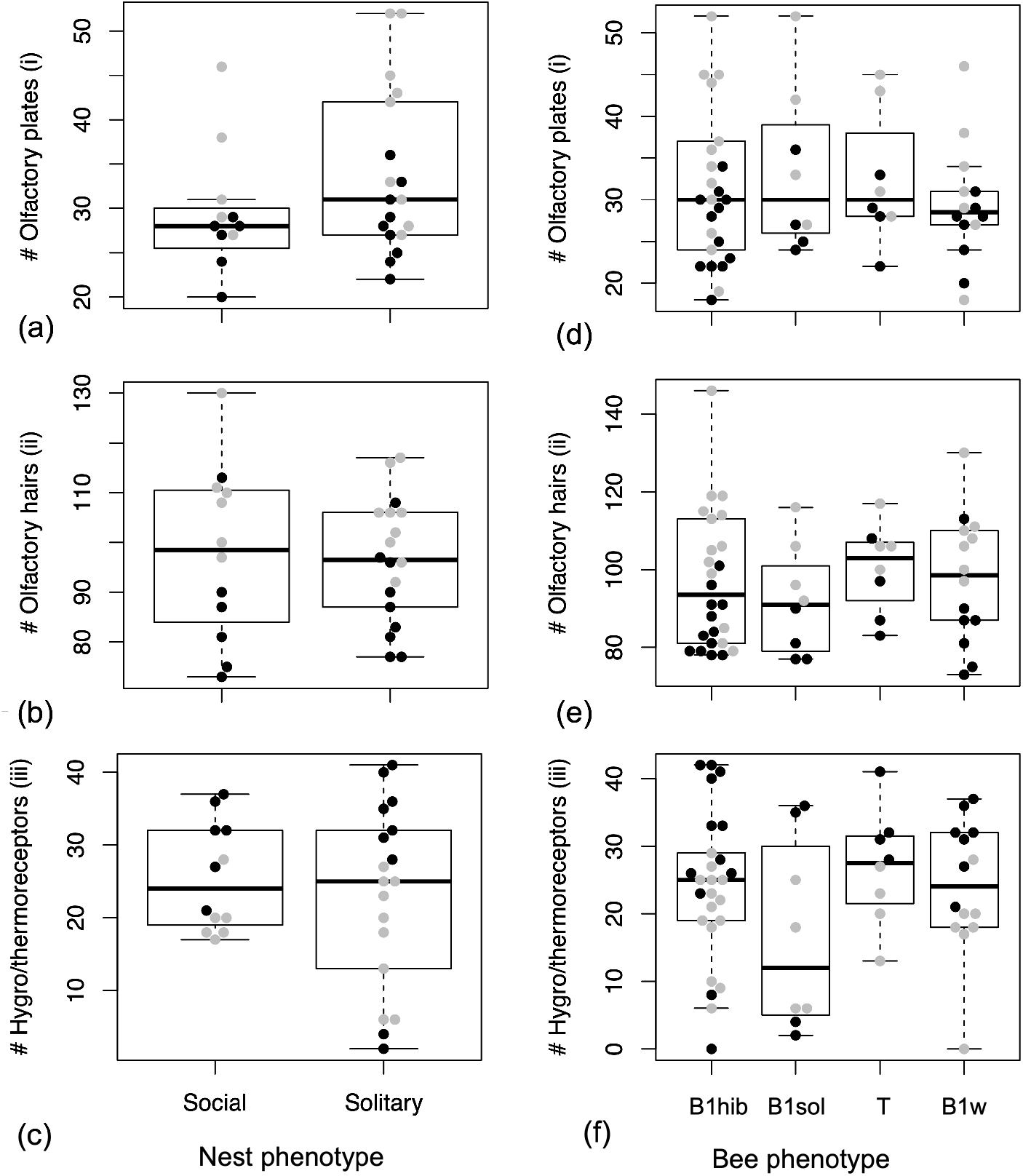
Numbers of (a) and (d) olfactory plates (top row); (b) and (e) olfactory hairs (middle row); and (c) and (f) hygro/thermoreceptive pores, (bottom row); for Halictus rubicundus from Scotland collected from (a-c) social and solitary nests and according to individual phenotype (d-f) after being transplanted to the south-east in 2020. For figures 4d-4f: B1hib female that emerged in the SE in 2020 but did not provision that year (will hibernate and provision in 2021); B1sol female that emerged in the SE in 2020 and provisioned a nest as a solitary foundress; T female that emerged in SCO in 2019, was transplanted to the south east and provisioned a nest as a solitary foundress in 2020; B1w female that emerged in the SE in 2020 and provisioned a social nest in the same year as a worker. Black points represent sensilla counts on segment 11 and grey points on segment 12. Box plots show median and quartiles across segments 11 and 12. There were no significant differences in counts of any sensilla type across the groups.

**Table 4 :**
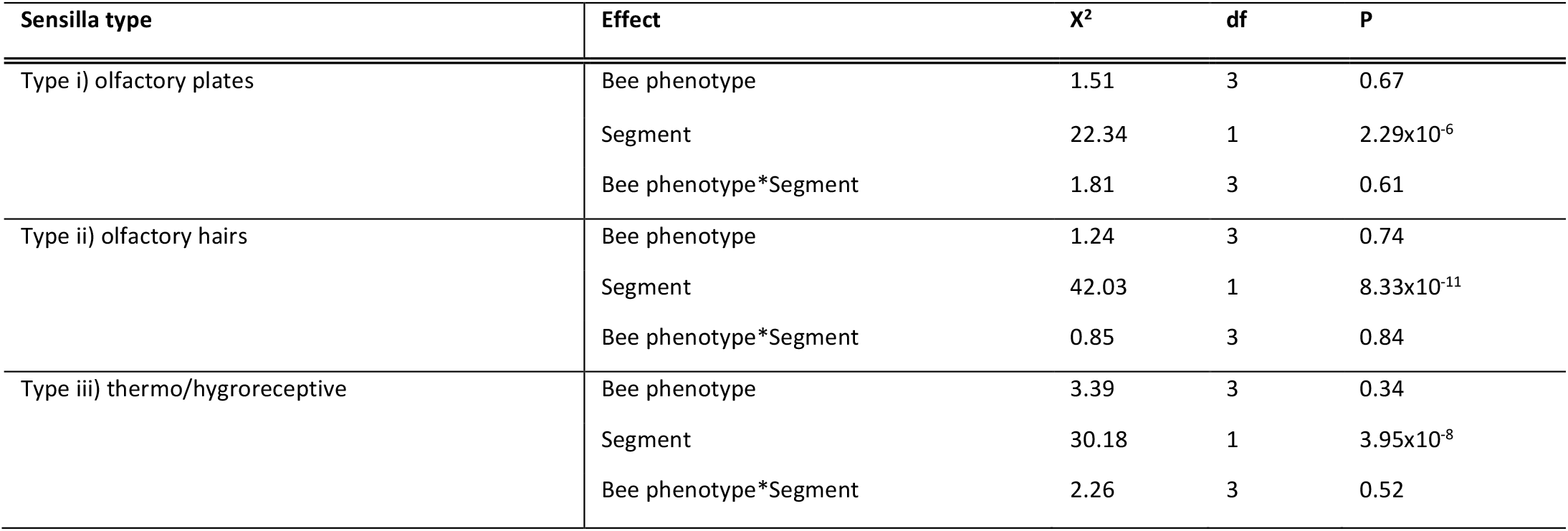
LMM results showing effect of bee phenotype (future reproductive/worker/solitary foundress 2019 and solitary foundress 2020) on antennal sensilla counts across segments (p-values for main effects are calculated independent from interaction effects which in all cases are non-significant).

## Discussion

The social Hymenoptera have contributed much to our understanding of the proximate and ultimate factors that underlie the evolution of sensory systems. Attention has focussed chiefly on how sociality has imposed selection to optimise olfactory communication, enhancing social cohesion through nest-mate recognition and supporting caste polyphenisms (Wcislo 1997; Ozaki et al. 2005; Spaethe et al. 2007; Renner and Nieh 2008; Riveros and Gronenberg 2010; Gill et al. 2013; Couto et al. 2017; Grüter et al. 2017; Elgar et al. 2018). In the current study we measured sensilla density in the socially plastic sweat bee *Halictus rubicundus*. We found that when bees were transplanted from Scotland, where they are solitary, to the south where they can be social, their offspring had higher densities of olfactory hairs. These results suggest that the density of olfactory hairs may, like sociality, be phenotypically plastic in *H. rubicundus*.

We also found evidence for between-region differences in the density of basiconic/trichoid olfactory hairs (and hygro/thermoreceptive sensilla) for untransplanted bees, with mid-latitude bees having higher densities than more northerly Scottish bees. Differences were not perfectly correlated with the expected degree of sociality, however, because bees from the warm south-west, which is where we expect sociality to be the dominant strategy did not differ from mid-latitude or Scottish bees. This contrasts with the results of Wittwer et al (2017) who showed that in another halictid, *Lasioglossum albipes*, females from social populations have higher densities of olfactory hairs than conspecifics from solitary populations. Unlike *H. rubicundus*, this species is not socially plastic, and fixed differences in social structure between populations may have resulted in canalising selection on antennal sensilla density in L. albipes but *not H. rubicundus*.

The results from our transplant experiment also suggest that in *H. rubicundus* olfactory hair density is not canalised and may be phenotypically plastic. Our results suggest that olfactory hair density may vary depending on the conditions that bees experience during development. When Scottish bees were transplanted to the south-east, their offspring that developed there (where air and soil temperatures in June and July are 2-5°C warmer than their native site; table S2) had greater densities of trichoid/basiconic hairs. It may be that in H. rubicunudus warmer temperatures during development, rather than social phenotype, lead to greater densities of olfactory hairs. A similar pattern is seen in the parasitoid wasp Trichogramma; when males develop under colder temperatures they have reduced numbers of basiconic hairs (Pinto et al. 1989). A direct response to temperature might explain why untransplanted foundresses from the south-west, where bees are often social, but temperatures are not as high as in the south-east, did not have higher basiconic hair densities than non-social populations (Fig. 3).

In addition to plasticity within a population, we found evidence that olfactory hair density (type ii, basiconic/trichoid sensilla) varies across regional populations in *H. rubicundus*. In contrast we saw no differences in the densities of placoid (plate-like, type i) olfactory sensilla across regions. This may relate to the function of these receptor types. Basiconic/trichoid hairs respond to contact with the CHCs of other individuals, and so are likely to be required for nest-mate recognition and communication, while placoid plates are thought to be involved in longer range olfaction to detect food and hosts (Ozaki et al. 2005; McKenzie et al. 2016; Couto et al. 2017; Pask et al. 2017). While food availability almost certainly varies across regions we know of no consistent regional variation in diet breadth in this species (although this remains to be rigorously studied), which fits with the pattern we see here.

Bees collected from Belfast at a mid-latitude had the highest density of olfactory hair-like sensilla (basiconic/trichoid; type ii), higher than bees from Scotland (450km north of Belfast) but statistically equivalent to bees collected in the south-west. We predicted that *H. rubicundus* collected further south would have the greatest densities of olfactory sensilla to support social communication between nestmates (Wittwer et al. 2017; Elgar et al. 2018). In the south, summers are typically warmer and longer and a higher proportion of the population is expected to exhibit social behaviour more regularly. While we did see regional differences in olfactory sensilla density, bees from the south-west did not have the highest densities as we would have predicted based on clines in sociality. The regional differences we saw may be related to developmental, potentially temperature-dependent plasticity, which could limit the scope for fixed population level differences to evolve in the predicted direction. Indeed, data from nearby weather stations lends support to the possibility that warmer temperatures during development might lead to higher densities of olfactory hair-like (basiconic) sensillae (see above; Pinto et al. 1989). The maximum and average air temperatures when the bees collected in Belfast were developing (in June and July) was up to 1°C higher than in the south-west and 1-2 °C higher than in Scotland (although the minimum temperatures in Belfast tend to be lower than in the south-west; see supplementary table S2 for complete climate data). Longitudinal studies of these same populations, reciprocal transplant experiments (from south to north) and more accurate measurements of soil temperature would help to elucidate the extent to which genetically fixed differences in olfactory hair density and plastic, temperature-dependent expression contribute to the population level differences we saw in this trait.

We also found that the density of sensilla (type iii) involved in the perception of humidity, temperature and CO2 varies across regions, but unlike olfactory hairs this does not appear to be plastic. Bees from a mid-latitude and the north of Scotland (Belfast and Scotland) have more thermo/hygro receptive sensilla than bees from the south-west (400 km south of Belfast and 850 km south of the Scottish site). The density of these sensilla does not appear to be plastic, however, as it was not influenced by where Scottish bees developed or their social phenotype. This pattern may be a result of a more extreme, variable climate in the north, which leads to consistent selection for higher densities of thermo/hygro receptors. Sweat bees including *H. rubicundus* are highly sensitive to temperature and rain. Flight activity is constrained by low ambient temperatures, lack of sunlight and rainfall (Potts and Wilmer 1997). In the north, bees experience colder ambient temperatures and greater rainfall. Bees with more of these receptors may be more sensitive to current and oncoming weather conditions, so that they have reduced mortality and improved foraging based on the climate. The potentially severe fitness consequences of incorrect perception of climatic cues may explain the lack of plasticity in the number of thermo/hygro receptors if selection has a strong and canalising effect on this trait.

Our results add a new dimension to the growing body of evidence that developmental temperature may contribute to adaptations which support sociality in the Hymenoptera. While previous studies suggest that temperatures experienced by developing larvae contribute to individual differences in social phenotype (i.e. caste polyphenism: Czekońska and Tofilski 2020; Becher et al. 2009; the development of status badges: Green et al. 2012; memory formation: Jones et al. 2005 and olfactory learning: Anton and Rossler 2021), sensilla density in the socially plastic *H. rubicundus* may be a more general response to the environment that is not directly related to the social phenotype of the nest or individual (i.e. a worker, solitary foundress or queen). Scottish bees that developed in the south-east had more olfactory hairs than their conspecifics that emerged in Scotland irrespective of their caste or nest phenotype.

In the Halictidae, Wittwer et al (2017) found that halictid bee species that had reverted back to solitary nesting from a state of sociality had reduced olfactory hair density compared to social species and ancestrally solitary species. They suggest that this is because dense olfactory sensilla are a pre-adaptation that facilitates the evolution of sociality and may contribute to the repeated evolutionary transitions to sociality seen in the halictid bees. Our results expand on this, suggesting that olfactory hair density may also be phenotypically plastic. More broadly, plasticity in traits such as this might contribute to the evolutionary lability of sociality in the Halictidae and in the Hymenoptera, acting alongside social plasticity in the hymenopteran ancestor to provide the ‘flexible stem’ which allowed for repeated evolutionary transitions to obligate sociality across the order (West-Eberhard 2003; Wittwer et al. 2017).

## Acknowledgements

Many thanks to Gavyn Rollinson and Sharon Uren for SEM training and technical support. Version 4 of this preprint has been peer-reviewed and recommended by Peer Community In Evolutionary Biology (https://doi.org/10.24072/pci.evolbiol.100140).

## Data, scripts and codes availability

Data andf code are available online: http://hdl.handle.net/11667/197.

## Supplementary material

Supplementary material are available online: http://hdl.handle.net/11667/197.

## Ethics

This research adheres to ethical codes of practice at the University of Exeter. No vertebrates were used in this research. Bee antenna samples were taken from living and dead stored specimens. Handling during antenna removal of live specimens was done as quickly as possible to minimize stress to individuals. Bees frequently lose parts of and whole antennae in nature and we did not see any negative effects on the survival or provisioning behaviour of live sampled individuals compared to intact bees. The sample sizes reported were considered appropriate to maximize statistical power while reducing the number of individuals involved in the experiments. Individuals were sampled from large populations > 500 individuals and numbers taken did not pose a threat to the persistence of these populations. The transplant experiment did risk causing gene flow (due to escape/mating) between Scottish and native bees in the south east (SE). We chose a transplant site with no records of H. rubicundus to minimize this risk. We have never found any H. rubicundus nesting nearby and the soil in the SE, which is dense clay, is unsuitable for nesting or hibernation of this species. At the end of the transplant experiment any surviving bees were taken back to their natal site in Scotland whilst they were hibernating in the buckets. We did not see any H. rubicundus at the site when we returned in the year following the transplant (2021). As such we believe that the risk of us causing gene flow between H. rubicundus populations is negligible.

