## Supplemental tables S1 and S2 for "Sensory plasticity in a socially plastic bee"

Table S1: Intraclass correlation coefficients for intra-rater reliability of all antenna types

| **Sensilla type** | **N** | **ICC** | **95% CI** | **F** | **p** |
|---|---|---|---|---|---|
| Type i) olfactory plates | 33 | 0.89 | 0.78 < ICC < 0.94 | 16.9 | $6.28\text{x}10^{-13}$ |
| Type ii) olfactory hairs | 33 | 0.85 | 0.72 < ICC < 0.93 | 12.3 | $1.02\text{x}10^{-10}$ |
| Type iii) thermo/hygroreceptors | 19 | 0.94 | 0.85 < ICC < 0.98 | 30.6 | $2.43\text{x}10^{-10}$ |

Table S2: Air temperature data taken from the wunderground website from nearest weather stations to each site where bees were collected. Shown are monthly maximum, minimum and mean temperatures (means ± standard deviations across all months) in years that bees that were collected from each site developed. Data from two years are provided for Migdale (Scotland) as bees were collected in two years (2018 and 2020 after developing in 2017 and 2019 (www.wunderground.com/history). Also included are soil temperatures recorded at depths of 1cm and 15cm at Migdale and Knepp sites in 2019 and 2020 respectively (using TinyTag data loggers; Gemini data loggers). At Knepp 1 logger was deployed and recovered at 1cm and 1 at 15cm. At Migdale 4 loggers were deployed and recovered at 1cm and 2 at 15cm.

| **Population** | **Nearest weather station** | **Year** | **Month** | **Max air temp (°C)** | | **Min air temp (°C)** | | **Average air temp (°C)** | | **Soil temp (1cm) ± SD (°C)** | | **Soli temp (15cm) ± SD (°C)** | |
|---|---|---|---|---|---|---|---|---|---|---|---|---|---|
| **Migdale (Scotland)** | Inverness airport | 2017 | April | 11.7 | 16 ± 3.0 | 3.5 | 6.9 ± 3.0 | 7.8 | 11.7 ± 2.8 | | | | |
| | | | May | 16.8 | | 5.2 | | 11.7 | | | | | |
| | | | June | 17 | | 9.0 | | 13.1 | | | | | |
| | | | July | 18.6 | | 9.9 | | 14.2 | | | | | |
| | | 2019 | April | 13.3 | 15.5 ± 3.1 | 3.2 | 6..9 ± 3.9 | 8.5 | 11.4 ± 3.4 | 11.3 ± 5.0 | 14.7 ± 6.3 | 10.3 ± 3.1 | 14.3 ± 3.8 |
| | | | May | 12.8 | | 4.2 | | 8.9 | | 13.3 ± 6.5 | | 13.0 ± 3.0 | |
| | | | June | 16.2 | | 8.2 | | 12.4 | | 15.6 ± 5.4 | | 15.2 ± 2.7 | |
| | | | July | 19.6 | | 11.8 | | 15.7 | | 18.4 ± 4.9 | | 17.8 ± 2.7 | |

| | | | | | | | | | | | | | |
|---|---|---|---|---|---|---|---|---|---|---|---|---|---|
| **Boscastle and Bodmin (south-west)** | Newquay airport | 2017 | April | 11.4 | 15.3 ± 2.9 | 6.6 | 10.9 ± 3.3 | 9.7 | 13.6 ± 3.0 | | | | |
| | | | May | 14.7 | | 10.0 | | 13 | | | | | |
| | | | June | 17.4 | | 13.0 | | 15.6 | | | | | |
| | | | July | 17.7 | | 14.0 | | 16.2 | | | | | |
| **Belfast (North)** | Belfast City aiport | 2019 | April | 11.9 | 15.6 ± 3.3 | 6.3 | 9.3 ± 3.2 | 9.0 | 12.4 ± 3.2 | | | | |
| | | | May | 14.1 | | 7.4 | | 10.9 | | | | | |
| | | | June | 16.9 | | 9.9 | | 13.4 | | | | | |
| | | | July | 19.5 | | 13.6 | | 16.4 | | | | | |
| **South-East (Knepp)** | Gatwick airport | 2020 | April | 17.2 | 19.8 ± 2.0 | 5.5 | 9.3 ± 3.3 | 11.4 | 14.7 ± 2.7 | 14.1 ± 4.5 | 18.1 ± 6.7 | 15.3 ± 2.5 | 18.3 ± 3.4 |
| | | | May | 19.3 | | 7.6 | | 13.7 | | 17.1 ± 5.8 | | 18.2 ± 2.8 | |
| | | | June | 20.7 | | 11.6 | | 16.2 | | 20.8 ± 7.5 | | 19.7 ± 3.0 | |
| | | | July | 21.9 | | 12.5 | | 17.5 | | 20.7 ± 6.1 | | 20.1 ± 3.0 | |
